# Hyaluronic acid hydrogels: Establishing a sustained delivery system for extracellular vesicles

**DOI:** 10.1101/2025.01.29.635521

**Authors:** Yashna Chabria, Joanne O’Dwyer, Emma McDermott, Peter Owens, Aoife J. Lowery, Garry P. Duffy, Róisín M. Dwyer

## Abstract

Extracellular vesicles (EVs) are versatile transporters of genetic cargo with enormous potential in the therapeutic setting. Scalable production of EVs, and routes to overcome rapid clearance are required. Biocompatible hydrogels may support precise, localized delivery of EVs to target sites. This study aimed to establish sustained production of EVs in a scalable 3D dynamic bioreactor and to fabricate hydrogels using tyramine-modified hyaluronic acid (HA-TA) to study EV integration and release patterns.

MDA-MB-231 cells transduced with lentiviral GFP fused with CD63, were cultured in a 20kD dynamic hollow fiber bioreactor and GFP-EVs harvested over five weeks. GFP-EVs were characterized by Nanoparticle Tracking Analysis(NTA), Western Blot(WB) and Transmission Electron Microscopy(TEM). Tyramine modified hyaluronic acid(HA-TA) hydrogels were formulated via enzymatic crosslinking using hydrogen peroxide and horseradish peroxidase, to investigate EV release patterns in static and dynamic conditions. Hydrogel swelling was recorded at 1-72 hrs and hydrogels were loaded with GFP-EVs to assess distribution and release by Scanning Electron Microscopy(SEM) and NTA respectively. GFP-EV uptake was assessed by confocal microscopy.

Longitudinal GFP expression was demonstrated in transduced cells and released EVs throughout bioreactor culture. TEM and NTA demonstrated successful isolation of EVs of 30-200 nm in size with intact lipid bilayers (average 4×10^9^ EVs/harvest). Initial harvests exhibited subpopulations of larger EVs, which disappeared upon serum withdrawal. WB verified the presence of EV markers CD63, TSG101, and CD81. HA-TA hydrogels were successfully formed and swelling assays revealed the requirement for higher concentrations of HA-TA and crosslinkers for scaffold stability and continued swelling. GFP-EVs were successfully incorporated into the hydrogels with variable release patterns observed over time, depending on EV concentration and hydrogel formulation. EV clusters in hydrogels were visualized by SEM. Investigation of GFP-EV release patterns under static and dynamic conditions highlighted a significant increase in release under fluid flow conditions. Efficient transfer of released EVs to recipient cells was also demonstrated *in vitro*.

The data demonstrate the potential for scalable production of engineered EVs in serum free conditions and subsequent incorporation into HA-TA hydrogels for sustained release. These biocompatible hydrogels hold promise for tuneable delivery of therapeutic EVs in a variety of disease settings.

## Introduction

Extracellular vesicles (EVs) are endogenous cellular nanoparticles that consist of a lipid bi-layer with transmembrane proteins and internal cargo (1,2). It is well established that cells release vesicles of different size, content and function. Small EVs (sEVs) are nano entities that are less than 200nm in size and have been recognized as key regulators of various pathological processes. sEVs are believed to be released by all cell types and are known to play a crucial role in intercellular communication by enabling efficient cargo transfer (3). EVs are loaded with peptides and nucleic acids that enable cellular crosstalk (4). Upon endocytosis EVs elicit alterations in the physiological state of recipient cells through a variety of mechanisms (2,3,5). These vesicles are believed to retain features of the secreting cell, making them potential alternatives for cell therapies.

The major limitation of cell therapies such as high costs, invasive harvesting procedures, extensive regulatory issues, and immune rejection can potentially be effectively overcome by EV therapeutics (6). The small size of EVs has benefits such as the ability to by-pass the blood brain barrier and capillary beds, potentially improving uptake at the target site.

The most common route of EV administration has been through systemic or localized doses that can result in rapid elimination. It has also been found to result in accumulation at non-target sites mainly the kidneys, liver and spleen (6). To ameliorate these effects, biomaterial-based systems are being tested to promote retention at the target site. Multifold materials both synthetic and natural such as hydrogels, scaffolds, sponges, electrospun fibers and many more are being investigated to enhance sEV bioavailability and reduce the requirement for multiple high doses to elicit substantial effects (7). Hydrogels exhibit excellent biocompatibility and are therefore one amongst the most extensively used biomaterials for a plethora of medical applications. The 3D hydrophilic polymeric matrices are classified into three categories based on their origin: natural, synthetic and hybrid gels. Due to a high affinity to water these 3D constructs are highly porous, making them ideal scaffolds for the delivery of therapeutics (3,5). The complex 3D lattices enable encapsulation and protection of hydrophilic molecules by preventing rapid degradation and clearance. Hydrogels are also effective sustained release systems as the encapsulated molecules are released based on the porosity and degradability of the crosslinked network (5,8) (Figure 1). Natural hydrogels have been used for several decades for tissue repair. Hyaluronic acid/hyaluronan (HA) is one such natural polysaccharide that is abundant in the body, mainly the extracellular matrix (ECM) (9,10). Native HA is rapidly turned over in the body due to the bioactivity of hyaluronidase, which can be attenuated through tyramine modification of the HA molecules thereby enabling sustained release of the therapy over the desired period of time (9,11). Hyaluronic acid hydrogels have previously been investigated for enabling effective release of particles in EV size range for tissue repair and regeneration (10–13). However, the EV distribution and release patterns are yet to be investigated. These delivery systems may improve therapeutic outcomes and reduce the dose of EVs required (5,6,14).

**Figure 1:**
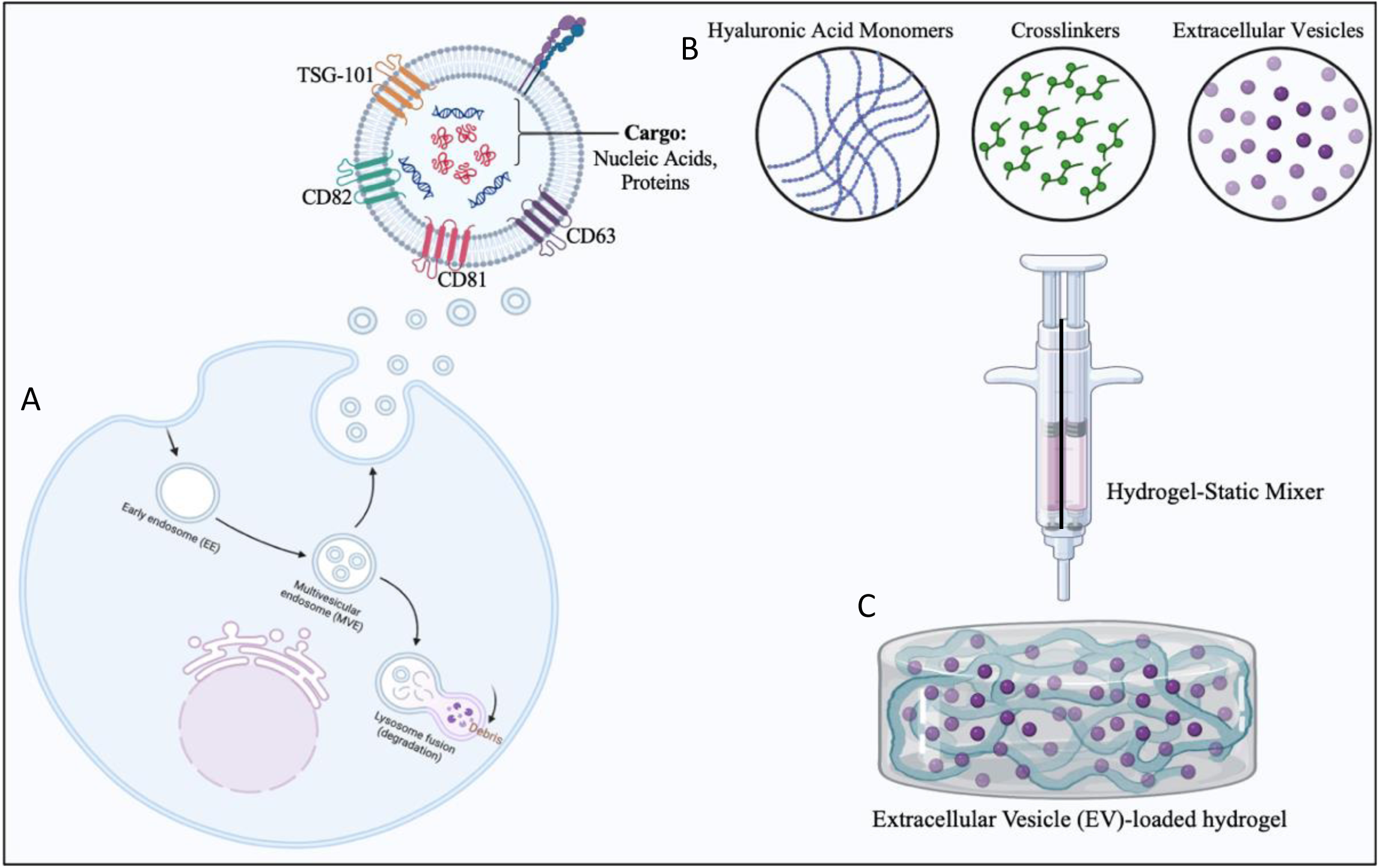
A. Schematic of EVs released from cells scaled up in a bioreactor system B. Formation of hyaluronic acid (HA) hydrogels by chemical crosslinking C. Fabrication of EVs loaded 3D tyramine modified HA hydrogels enabling sustained release (created using Biorender).

Also relating to the dose, insufficient yields of EVs obtained from traditional 2D cell cultures have prompted researchers to investigate methods for improving EV production. Scalable, reproducible EV production is essential for replacing cell-based therapies with EV therapeutics (15). These platforms include roller bottles and microfiber-bed systems which augment cell loading and attachment by increasing surface area, and 3D culture systems such as hollow fiber bioreactors or microcarriers which facilitate inoculation of high cell numbers in a compact space (16).

The aim of this study was to produce EVs in a scalable bioreactor system, incorporate these into tyramine-modified hyaluronic acid (HA-TA) hydrogels and analyse the EV release profiles in static and dynamic conditions (*Figure 1*).

## Materials and Methods

### Lentiviral transduction of MDA-MB-231 cells

MDA-MB-231 triple negative breast cancer cells (LGC Limited, UK) stably transduced with Exosome Cyto-Tracer, pCT-CD63-GFP (System Biosciences) were used for EV production. The cells were routinely cultured in DMEM-Glutamax media containing 10% foetal bovine serum (FBS) and 100LIU/ml penicillin/100 µg/ml streptomycin (Pen/Strep). The lentiviral construct consisted of a GFP-fusion protein, hybrid RSV-5’LTR promoter, cytomegalovirus (CMV) promoter and a puromycin resistance gene. The cyto-tracer fuses the Green Fluorescent Protein tag to CD63 present on the plasma membrane resulting in GFP expression in the cells and the released sEVs. Cells were transduced at a multiplicity of infection (MOI) of 10 in serum free media with 2 µg/ml Hexadimethrine bromide (Polybrene, Sigma-Aldrich) for 6 hours. The cells were then maintained in fresh complete media for 48 hours followed by selection of transduced cells in the presence of puromycin (4 µg/ml) (Sigma-Aldrich) over the next 14 days. GFP expression in live cells was confirmed by fluorescence imaging using EVOS FL Cell Imaging System (Thermo Fisher Scientific) and flow cytometry (Cytek Northernlights NL2000) and analysed by Flow Jo^TM^ (Becton Dickinson & Company (BD), version-10.8.1). Successful GFP transfer to EVs was confirmed by EV uptake studies using confocal microscopy (Olympus Fluoview 3000).

### Scalable EV production in bioreactor system

A Fibrecell bioreactor system was employed for EV production. A medium sized cartridge with a cut-off of 20kDa (C2011, Fibercell Systems; Frederick, MD, USA) was used for the culture of GFP MDA-MB-231 (MDA-GFP) cells that enabled the retention of vesicles (40– 1000nm in diameter, molecular weight >500 kDa) and the transfer of nutrients and waste-products through the fibers. Following the manufacturer’s guidelines the bioreactor cartridge was pre-treated in preparation for inoculation with 1 x 10^8^ cells. Cells were first accustomed to the 3D culture system in standard growth media (DMEM-Glutamax +Pen/Strep (1% P/S) + 10% FBS). Glucose consumption was monitored using glucose strips (GlucCell Glucose Strip, GC001001, KDBIO) attached to a glucometer (GlucCell Meter, GC001000, KDBIO). Following manufacturer’s guidelines when the glucose consumption rates peaked, the media was then switched to serum free media (DMEM + 10% Fibercell Systems chemically defined medium for high density cell culture {CDM-HD; Fibercell Systems, Frederick, MD, USA} protein-free supplement + Pen/Strep). Cells were cultured in the system for 52 days resulting in 24 EV harvests (20ml total volume collected each time). The EV rich conditioned media was harvested 3-5 times/week based on glucose consumption rates.

### EV Isolation by Size Exclusion Chromatography (SEC)

The EV-rich media harvested from the FiberCell underwent differential centrifugation at 300 x g and 2000 x g for 10 minutes each, followed by filtration (0.22µm filter) to eliminate cell debris and large proteins. The qEV 10 columns (35mm series-350nm cut-off) and the Automated Fraction Collector (AFC) (both from Izon Science, Lyon France) were used for isolating GFP-sEVs from cell culture media. EV isolation was performed following manufacturer guidelines. Briefly, prior to loading of samples the column was primed with sterile filtered PBS. Following this, the required parameters were set up on the AFC. These included fraction number, fraction size and void size for collection. The AFC first eluted out the void volume after which 7 x 5ml fractions of sEVs were collected. sEV fractions were stored at −80° for subsequent analysis.

### Characterization of EVs

#### Nanoparticle Tracking Analysis (NTA)

The size distribution and concentration of isolated sEVs were assessed and characterized by Nanoparticle Tracking Analysis (NTA) (Nanosight NS300, Malvern Panalytical, UK) using a 405 nm laser source and EMCCD camera, running NTA software version 3.2 using optimised and validated protocols (17). EV samples were diluted in particle-free PBS, confirmed by NTA with fewer than 3 particles per frame observed. Daily calibration of the instrument was conducted using 100 nm polystyrene latex nanoparticles from Malvern Panalytical. For all samples the screen gain was maintained at 1 and the camera level was adjusted based on the sample visibility. The same detection threshold was maintained for all the fractions of the sample. A total of 3-5, sixty second videos were recorded for each sample to determine sEV size distribution and concentration. The concentration of total EVs per microliter was calculated, along with the specific count of those sized between 30 and 150 nm, indicative of the small EV (sEV) fraction.

#### Western Blot

EV protein suspended in Triton-X lysis buffer was assessed by western blot analysis to confirm expression of EV associated proteins including tetraspanins (CD 63, CD81, CD82), EV biogenesis marker TSG-101 and absence of endoplasmic marker calnexin (Table 1). Following manufacturer instructions, protein concentrations were first assessed using microBCA Assay (Pierce™, Thermo Fisher Scientific). Protein samples were then denatured in DTT (0.5 M) for 10 minutes at 70°C, loaded into the wells of a pre-cast 4-15% Mini-PROTEAN^®^ TGX™ Gel (Bio-Rad) along with a protein molecular weight standard (20–220 kDa) and run at 100 V for 1 hr. The proteins were then transferred to a nitrocellulose membrane on a transfer unit at 25V for 60 minutes. The blots were then blocked in 5% milk in TBS-T washing buffer for 2 hours and then probes for specific antibodies were applied as illustrated in Table 1. Following wash steps in TBS-T the blots were then incubated with respective secondary antibodies, followed by further wash steps and incubation with a chemiluminescent substrate solution (Clarity™ Western ECL-Bio-Rad Laboratories) to support imaging on Gel Doc™ XR+ and ChemiDoc™ XRS + Systems with Image Lab™ Software (Bio-Rad, version 5.2.1).

**Table 1:** Antibody targets for Western Blot (All antibodies were purchased from Abcam)

| *Antibody* | *ID* | *Dilution (Time)* | *Secondary* | *ID* | *Dilution (Time)* |
| --- | --- | --- | --- | --- | --- |
| CD 63 | Ab59479 | 1/1000 (16hrs) | Rabbit Anti-Mouse | Ab6728 | 1/5000 (1.5hrs) |
| CD 82 | Ab66400 | 1/1000 (16hrs) | Goat Anti-Rabbit | Ab66400 | 1/3000 (1.5hrs) |
| TSG-101 | Ab125011 | 1/1000 (16hrs) | Goat Anti-Rabbit | Ab30871 | 1/3000 (1.5hrs) |
| CD 81 | Ab79559 | 1/10,000 (1.5hrs) | Rabbit Anti-Mouse | Ab6728 | 1/10,000 (1.5hrs) |
| Calnexin | Ab22595 | 1/1000 (16hrs) | Goat Anti-Rabbit | Ab10286 | 1/3000 (1.5hrs) |

#### Transmission Electron Microscopy (TEM)

As previously described (18), sEV samples were first treated with primary fixative (2% Glutaraldehyde, 2% Paraformaldehyde in a 0.1 M Sodium Cacodylate/HCL buffer at a pH of 7.2). This step was followed by a secondary fixation step in 1% Osmium Tetroxide. The sEV pellet was finally dehydrated and embedded at 65^◦^C in resin before sectioning. 80– 100 nm thickness sections were taken using an ultramicrotome (Reichert-Jung Ultracut E). Sections were stained with 1.5% aqueous uranyl acetate followed by lead citrate and loaded onto a copper grid for imaging (Hitcachi H7000 Transmission Electron Microscope).

#### Concentration of EVs

For EV release studies, EVs were concentrated using Amicon® Ultra-15 centrifugal filter unit (Merck Millipore) following the manufacturers guidelines. Briefly, three 5ml fractions from the fibercell harvests were pooled together into the concentration column. The concentration unit was then spun in a swing bucket rotor centrifuge at 4000 x g for 20 minutes at 4°C. The EVs were then collected from the V-shaped concentration chamber. Success of concentration was determined by analysing final EV concentrations by NTA.

#### EV Uptake studies by Confocal Microscopy

EVs derived from MDA-GFP cells cultured in the bioreactor were used for uptake studies. Recipient wild-type (WT) MDA-B-231 cells were seeded on to glass slide chambers [µ-Slide 8 Well ^high^ (Ibidi-80806)] at a density of 5 x 10^3^ cells/well. Isolated GFP-EVs were applied at a concentration of 1×10^9^/well and incubated overnight. Prior to imaging the cells were fixed with 4% Paraformaldehyde and cell nuclei were counterstained with 4’,6-diamidino-2-phenylindole (DAPI) (Invitrogen). The cells were imaged using a laser confocal microscope (Olympus FluoView 1000) by taking multiple immunofluorescent Z-stack images at 60X magnification. Further image processing was performed using ImageJ software (19).

#### Fabrication of hyaluronic acid hydrogels for EV encapsulation

Tyramine modified Hyaluronic Acid (HA-TA) (250–350 kDa, 1-2% w/w tyramine substitution) was purchased from Contipro (Czechia) in a lyophilized form that was reconstituted in PBS (1% and 2% concentrations) overnight on a roller plate to ensure thorough wetting of the freeze-dried HA-TA powder. Gels were fabricated by chemical crosslinking using Hydrogen Peroxide (H_2_O_2_) and horseradish peroxidase (HRP) (final concentrations of crosslinkers are given below). As native H_2_O_2_ is toxic to EVs, HA-TA was mixed with both the crosslinkers separately and gelation was established using a benchtop hydrogel mixer (BHM) that consisted of a static mixer enabling rapid interactions between the two solutions to form intact hydrogels. Consistency of hydrogel size and shape was achieved by injecting the HA-TA dispersions, through the double syringe system, into a 3D cylindrical mould. Hydrogels were prepared using HA-TA at a concentration of 1% or 2% w/v. (Table 2). Two crosslinking densities were used for each HA-TA concentration to assess the effect of hydrogel mechanical properties and crosslinking on swelling and EV release.

**Table 2.** : **HA-TA Hydrogel formulations (HA-TA and crosslinker concentrations)**

| Formulation | HA-TA concentration | H<sub>2</sub>O<sub>2</sub> concentration | HRP concentration |
| --- | --- | --- | --- |
| 1% 1x | 1% w/v | 0.88 U/ml | 0.12 U/ml |
| 1% 2x | 1% w/v | 1.76 U/ml | 0.24 U/ml |
| 2% 1x | 2% w/v | 1.76 U/ml | 0.24 U/ml |
| 2% 2x | 2% w/v | 3.52 U/ml | 0.48 U/ml |

#### Hydrogel Swelling Assay

Hydrogel weight was measured immediately following crosslinking and prior to immersion in any liquid (m_0_), to analyse the swelling properties of the hydrogels. Hydrogel discs formed in the 3D moulds were immersed in 1ml PBS or DMEM in each well of a 24 well plate. At each time point (1-6 hours, 24, 48, 72, and 96 hours), the hydrogels were removed from the liquid, blotted dry, weighed (mt), and the swelling percentage was calculated using the following formula (20).

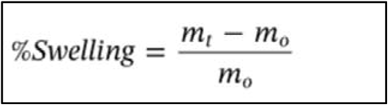

Hydrogel swelling is mainly determined by the strength of the crosslinks and the pH of the media/buffer. Swelling can provide an indication of the stability and release profile of the hydrogel.

#### SEM of EV distribution in hydrogel

Following initial processing of freeze drying (Telstar freeze-dryer) at the temperature of −59 degrees Celsius and the vacuum at 0.046 mBar for 18 hours, the internal morphology and arrangement of the hydrogel, along with the distribution of EVs within it, were examined using Scanning Electron Microscopy (Hitachi-S2600N). To prepare for imaging, samples were affixed onto aluminium stubs and coated with a thin layer of gold using a Gold Sputter Coater (Quorum Q150R ES plus) for 2 minutes. Additionally, the gels were embedded in resin to facilitate further investigation of EV structure within the hydrogel using Transmission Electron Microscopy (Hitachi-7500).

#### Static EV release Studies

GFP-sEVs were incorporated into the hydrogels using a static gel mixer. For this study, 1.5×10^11^ sEVs were loaded and all experiments carried out in triplicate. The sEV-loaded hydrogels were immersed in 1ml of PBS/well in a 24-well plate. The PBS was removed at each time point of the experiment and stored −80° C. Samples containing sEVs released from the hydrogels were then analysed using NTA.

#### Dynamic EV release Studies

To analyse EV release patterns from the hydrogels in dynamic conditions, the Single-Flow MIVO^®^ device (React4Life) was established with the aim to recapitulate the complexity of a 3D, dynamic culture system at the flow rate of blood flow *in-vivo*. Briefly, GFP-sEVs loaded in 2% 1x HA-TA hydrogels were placed in 0.8 µm transwell insert and loaded onto the MIVO chamber. The chamber was then connected via tubing to form a closed circuit system attached onto a peristaltic pump set at the flow rate of 17 rpm following manufacturer’s recommendation to mimic *in-vivo* blood flow rate. The MIVO circuit was then filled with sterile particle free PBS that was sampled out at different time points to analyse sEVs release patterns in dynamic conditions over time.

#### Statistical Analysis

When feasible, data is displayed as Mean ± SEM. Analysis was conducted utilizing Excel 2016 for Windows. A two-sample t-test was employed to compare swelling profile and the static and dynamic sEVs release patterns. Results with a p-value < 0.05 were deemed statistically significant.

## Results

### Confirmation of transduction and expansion in the bioreactor system

Successful lentiviral transduction of MDA-MB-231 cells with CD63-GFP was confirmed by fluorescence microscopy and flow cytometry analysis. Following puromycin selection, cells were visualized to be positive for GFP expression using fluorescence microscopy (Figure 2 (i) A) and flow cytometry (Figure 2 (i) B). MDA-GFP cells were successfully inoculated into the 3D fibercell bioreactor system to allow for a continuous harvest of sEVs for 33 days once stable culture was established based on the monitored glucose measurements. Throughout the 33-day period of EV harvest, a total of 24 x 20ml harvests of sEVs-rich cell-conditioned medium (CCM) were successfully collected for subsequent sEV isolation and characterization. The timeline of continuous culture and interventions in bioreactor is shown in Figure 2 (ii). Maintaining cell viability throughout sEV production was crucial to ensure production of high-quality sEVs with desired physio-chemical properties. Throughout the entire 52-day cell incubation period within the hollow-fiber bioreactor system, continuous glucose consumption was observed (Figure 2 (iii)). Initially, cells were maintained in DMEM supplemented with FBS for 14 days to support continuous cell proliferation and homeostasis in the 3D system. A noteworthy reduction in glucose rate was observed upon switching to serum-free conditions at day 15 until cells adapted to the nutrient starved conditions. However, once homeostasis was achieved, consumption rates consistently rose, confirming sustained cell bioactivity in the FiberCell bioreactor system in serum free conditions. The presence of hollow fibers provided ample surface area for cell attachment and growth, resulting in increased cell density over time. Due to high cell proliferation indicated by a glucose consumption rate of ≥1500 mg /day, cell numbers were reduced in line with manufacturer’s guidelines. A high glucose harvest was conducted on day 43, resulting in a rapid decline in glucose consumption rates. Glucose consumption rates then rapidly increased again over the subsequent days. Cells from the bioreactor system were also harvested at various time points (LGH1-day 21 and LGH 5-day 27) throughout the 52-day culture period along with sEVs to verify the maintenance of GFP expression. Robust GFP expression was validated in these harvested cells using flow cytometry (Figure 2 (iv)).

**Figure 2:**
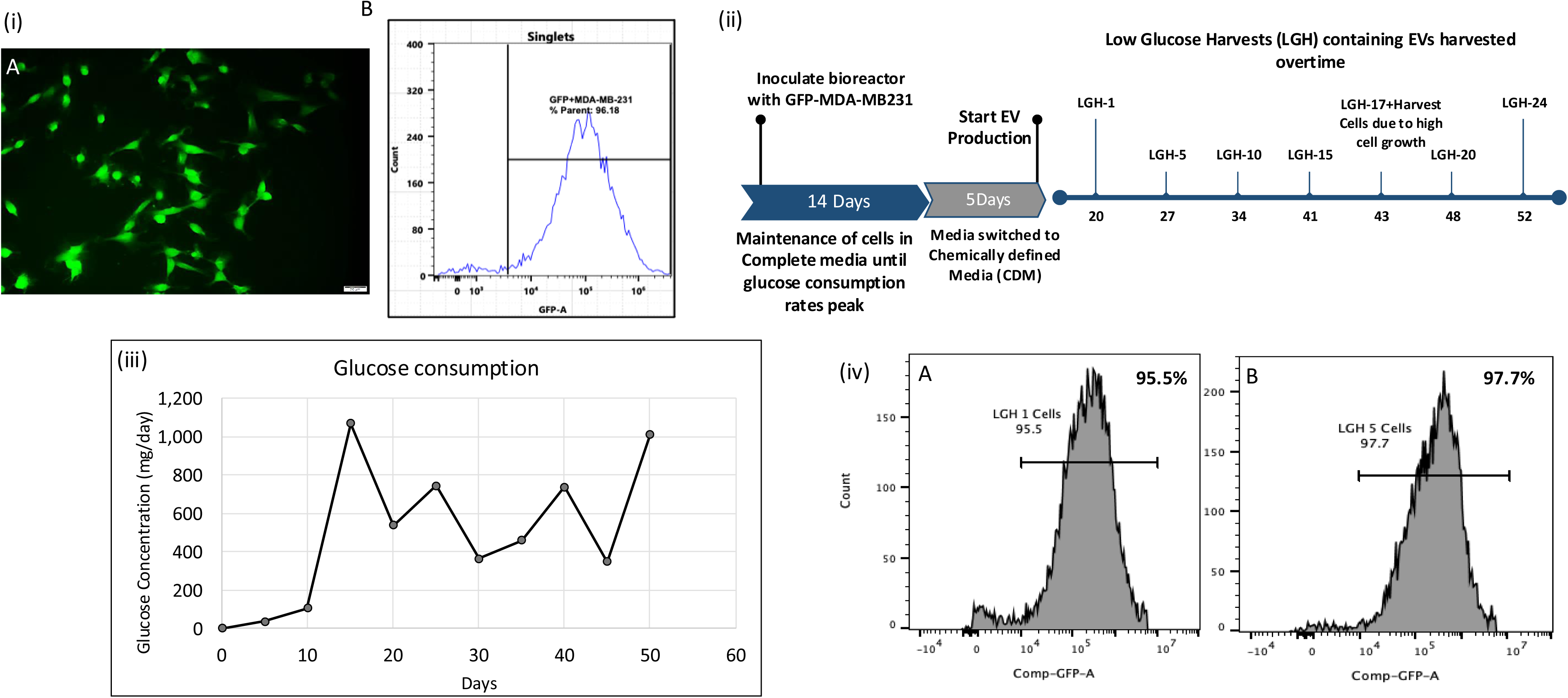
(i) Confirmation of GFP Expression in transduced MDA-MB-231 cells (A) Representative fluorescence imaging of GFP+ve cells (20x Magnification, scale bar 20µm) (B) Flow cytometry analysis confirming GFP expression (ii) Fibercell bioreactor workflow involved in GFP-EVs production (iii) Glucose consumption of cells maintained in the bioreactor over the 52-day time-period (iv) Confirmed GFP expression by flow cytometry analysis in cells harvested from the bioreactor system on (A) day 20 (LGH-1) and (B) day 27 (LGH-5).

### Characterization of GFP-sEVs harvested from the bioreactor system

From all the 24 (20ml) harvests of CCM, sEVs were successfully isolated by SEC that yielded 7 fractions/harvest. NTA analysis revealed EV concentration and size distribution of all the isolated fractions. A representative NTA graph shown in Figure 3 (i) exhibits distinct peaks indicating particle size range predominantly below 200nm. NTA also demonstrated procurement of high sEV yields in the range of 1.36 x 10^10^ to 1.46 ×10^11^ particles/ml with the average sEVs size of 154.6 nm. The transition from serum-containing media to CDM-HD serum-free media was reflected in the NTA analysis of the harvested sEVs. The red peaks observed on the NTA graphs (Figure 3 (i and ii)) distinctly differentiate early EV harvests from later ones. Early harvests exhibited a broad spectrum of irregular peaks with notably broader size distribution, indicating the gradual replacement of serum-containing media (Figure 3 (ii-A)). This transition is clearly illustrated by the clean peaks and consistent size distribution within the target range observed in subsequent EV harvests (Figure 3(ii-B)).

**Figure 3:**
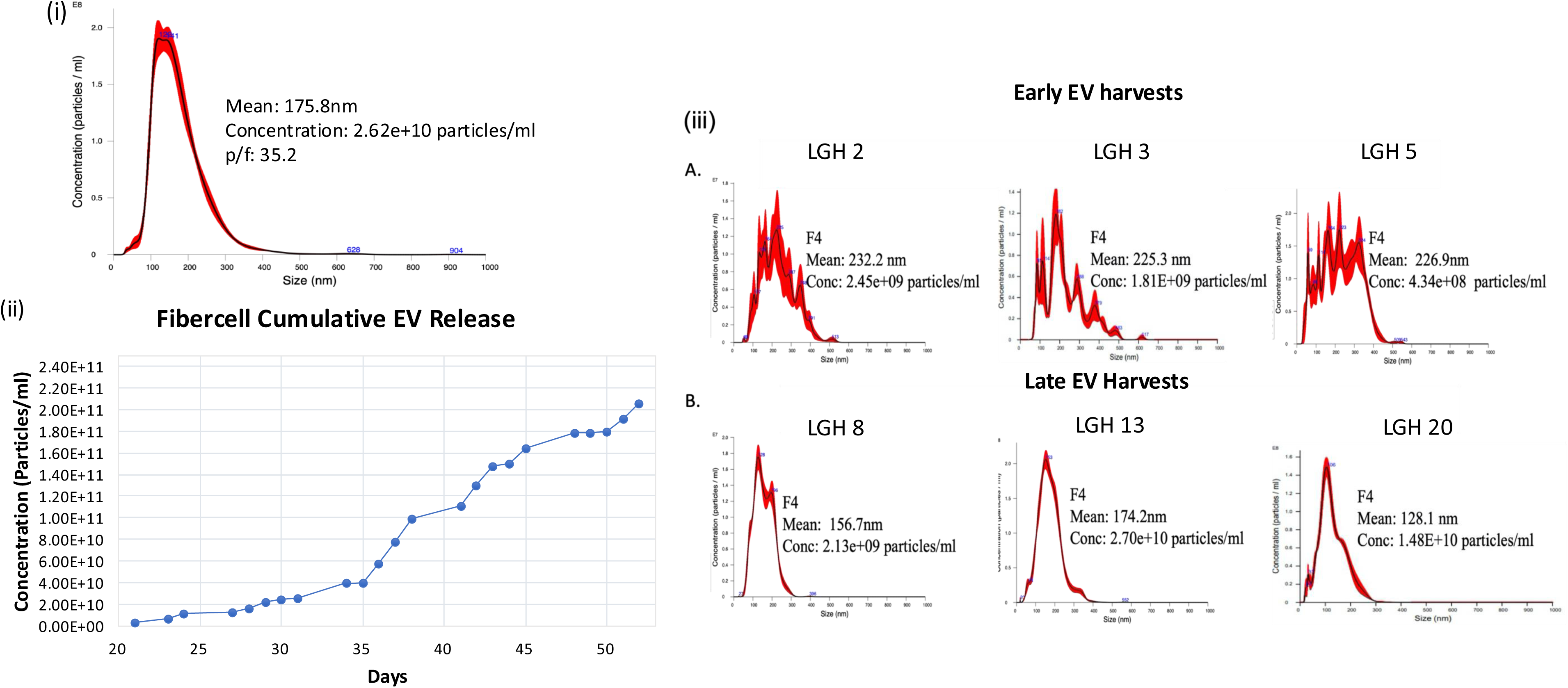
Characterization of EVs harvested from the FiberCell (i) Representative NTA graph demonstrating average EV mean size and concentration (EVs/ml) (ii) Cumulative EV graph of EVs procured from 24 LGH’s harvested over a period of 52 days (iii) Representative NTA graphs demonstrating (A) serum contamination in early EV harvests LGH 2, LGH 3 and LGH 5 compared to (B) later EV harvests LGH 8,13 and 20 obtained post switch to serum-replacement CDM media.

Consistent EV harvests were performed starting from day 20 to day 52. The cumulative yield demonstrated a steady rise in EV concentrations throughout this period (Figure 3 (iii)). By the end of 52 days, the total sEV yield reached 1.40 x 10^12^ sEVs/ml. To further assess the impact of concentrating sEVs rich fractions with concentration columns, sEVs size distribution and concentration of final sEVs concentrate (fraction 3-5) was assessed by tracking analysis (Supplementary Figure 1).

Electron microscopy imaging further confirmed the characteristic lipid bi-layer and desired size range of the isolated sEVs (Figure 4(i)). Upon estimation of protein content of all 7 fractions of sEVs samples, a consistent increase was observed in later fractions 6 and 7. sEV harvests from early (LGH 5), interim (LGH 10 and LGH 15) and later (LGH 20) timepoints were compared and a similar gradient of increasing protein levels was observed in fractions 6 and 7 (Figure 4 (ii)). Analysis of sEV protein revealed expression of EV associated proteins-tetraspanins (CD 63, CD 81), TSG-101 sEV biogenesis marker and absence of endoplasmic marker calnexin (Figure 4 (iii)). Finally, sEVs harvested from the FiberCell were analysed for GFP expression by performing uptake assays with wild type MDA-MB-231 cells by confocal microscopy. Following overnight incubation, GFP-sEV clusters were observed associated with the DAPI stained nuclei of cells (Figure 4 (iv)).

**Figure 4:**
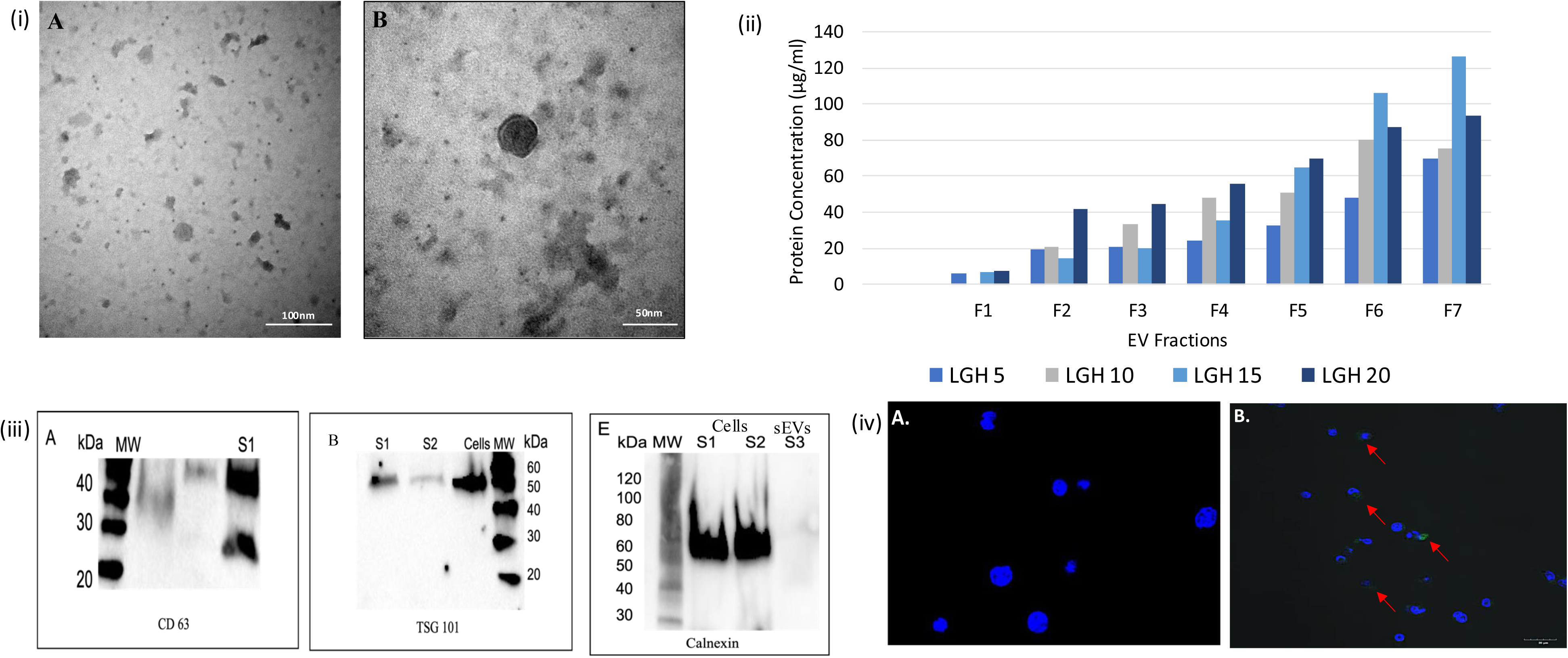
Characterization of EVs harvested from the bioreactor system (i) TEM imaging demonstrating characteristic lipid bi-layer (A) 120,000x (B) 200,000x (ii) Protein Concentrations of EVs obtained from early, intermediate and late EV harvests (Fractions 1-7) (iii) Western blot analysis of GFP-sEVs positive for tetraspanin CD63 (sEVs Sample 1), ESCRT pathway marker TSG 101 (sEVs sample S1 and S2 and cells; higher expression observed in cells) and negative for endoplasmic marker calnexin in (sEVs sample S3) and observed in cell samples (S1 and S2) (iv) GFP expression observed in EV uptake studies by confocal imaging (A) Control wild type MDA-MB231 cells stained with DAPI co cultured with no EVs (60X Magnification) (B) Wild type MDA-MB231 cells stained with DAPI co cultured with GFP EVs (60X Magnification), red arrows showing uptake of GFP EV clusters.

### HA-TA hydrogel formation and Characterization

To determine the optimal HA-TA concentration for vesicle release over time, four combinations of HA-TA hydrogel (1% 1x and 2x; 2% 1x and 2x) were established and analyzed. Hydrogels were formed using the benchtop mixer and 3D printed moulds. Previous literature indicates that hydrogel formation occurs rapidly post-injection through the benchtop mixer (11). Hydrogels were removed from the 3D moulds for imaging and further analysis. SEM imaging of the freeze-dried HA-TA (Figure 5 (i)), revealed successful formation of fibrillar structures demonstrating a mesh like network that is characteristic of hydrogels.

**Figure 5:**
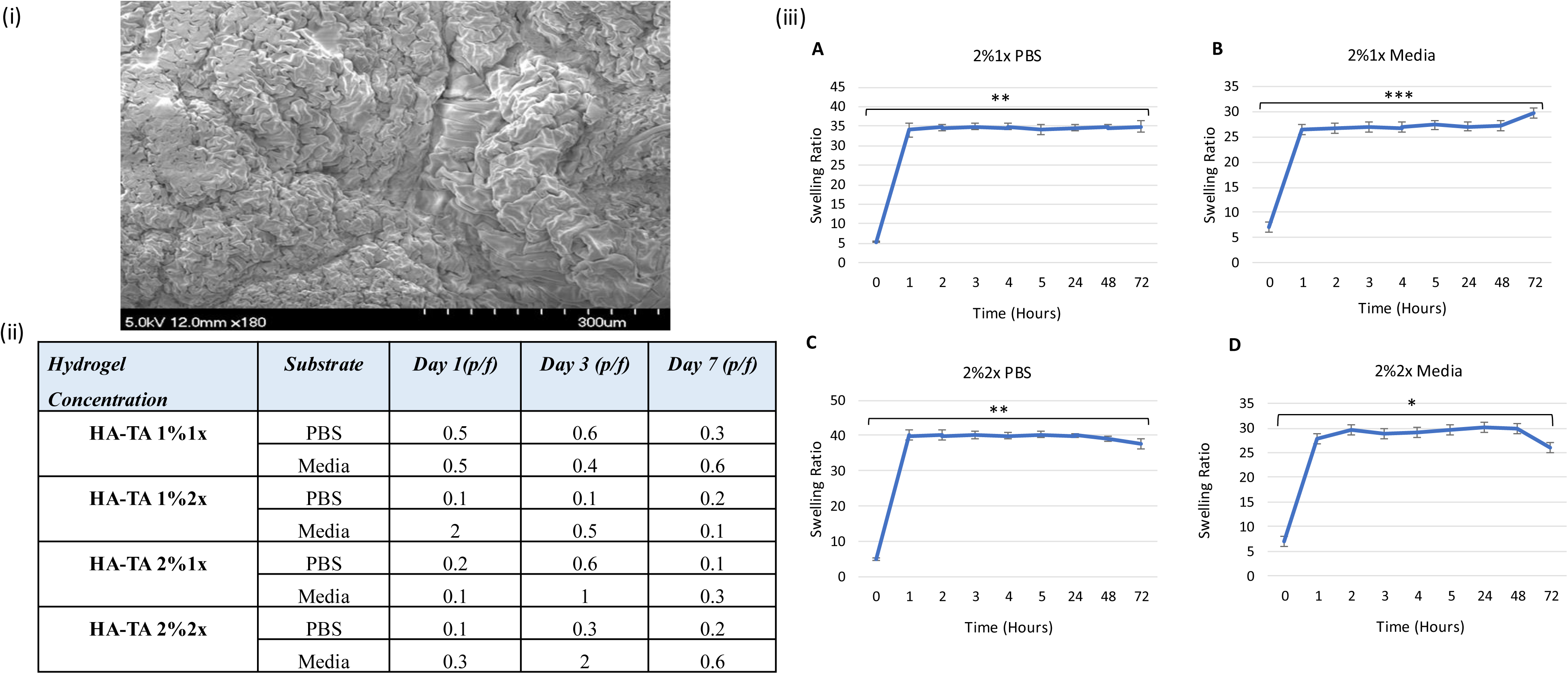
Characterization of HA-TA Hydrogels (i) Scanning electron microscopy images of HA-TA hydrogels taken within 1 hour of formation of gels, highlighting its characteristic fibrillar network under magnifications of 180x (ii) NTA particles/frame data of blank gels of varying HA and crosslinker concentrations highlighting undetectable levels of HA-TA particles in the EV size range over a period of 7 days (iii) Swelling assays demonstrating maintenance sustained swelling up to 72-hour time point in 2% hydrogels A. 2%1x (**P=0.002) in PBS B. 2%1x in Media 1x (***P=0.0004) C. 2%2x in PBS 1x (**P=0.002) D. 2%2x in Media1x (*P=0.03), Graph shows mean ± SEM (n=3).

To identify the most suitable HA-TA formulation for optimal EV release, all 4 formulations were analyzed for degradation (in the absence of EVs). The degradation products of the hydrogels were analyzed via NTA to observe if the hydrogel degradation would produce particles in the size range of the sEVs, which could impact the reliability of detected EV release patterns. Blank hydrogels underwent evaluation in both PBS and EV-depleted media conditions for a duration of 7 days. To ensure accuracy, the particle/frame (p/f) values in the NTA readings are required to fall within the detection threshold range of 10-50 (p/f). Across all hydrogel formulations, none to negligible release of particles within the size range of sEVs was observed (Figure 5 (ii)). The particles/frame parameter averaged between 0.3 and 0.6 particles/frame for 1% hydrogels in PBS and media, respectively.

Further analysis of the swelling behavior of all HA-TA hydrogel formulations was performed to identify optimal hydrogel formulations for EV release studies. Considering the influence of ionic strength on swelling, tests were conducted in both PBS and complete growth media. This criterion was crucial in selecting the most suitable hydrogel, ensuring stability for sustained EV release. Additionally, the assay assessed hydrogel integrity and shape retention. The 1% hydrogel formulations (1x and 2x) exhibited no swelling in either PBS or media, indicating rapid degradation and fragility, leading to their exclusion from further study (Supplementary Table 1).

In contrast, swelling profiles of 2% hydrogel formulations when monitored over 72 hours, revealed significant swelling within the initial hour followed by sustained weight retention in both PBS and cell culture media over time. Comparing the 2% 1x and 2x gels, the 2% 1x hydrogels exhibited consistent and reproducible swelling patterns over a longer duration (Figure 5 (iii A-D)). Conversely, while the 2% 2x gels displayed continuous swelling, a notable decrease in swelling ratio was observed at 72 hours in the media group (Figure 5 (iii B and D)). Consequently, the 2% 1x hydrogels were chosen for further investigation of EV loading and release as they demonstrated consistent swelling over time in both PBS and media (Figure 5 (iii)).

sEVs were effectively encapsulated into 2% 1x HA-TA hydrogels and subjected to subsequent imaging. SEM revealed the distribution of sEV clusters within the folds of the hydrogel’s crosslinked structure (Figure 6 (i)). Enhanced magnification illustrated the uniform distribution of sEVs apparently adhered to the inner surface of the 2% 1x HA-TA hydrogel. Following successful observation of sEVs released from EV-loaded hydrogels (supplementary figure) further investigations were carried out to assess the influence of dynamic flow conditions on GFP-sEV release patterns (n=3). Alternate hydrogels were exposed to either static or dynamic conditions, simulating *in vivo* blood flow, and the released sEVs were analyzed via NTA. sEV release under dynamic conditions was significantly increased compared to the static setup ((p=0.01) Figure 6 (ii)). An initial burst release of sEVs was observed on day 1, and overall EV release declined by day 7 after which further time points weren’t analyzed as it was beyond the detection threshold of the NTA. The average sEV release rate in the dynamic system remained at 3.4×10^7^ sEVs/ml, whereas in static conditions, it dropped to 5.3 x 10^6^ sEVs/ml.

**Figure 6:**
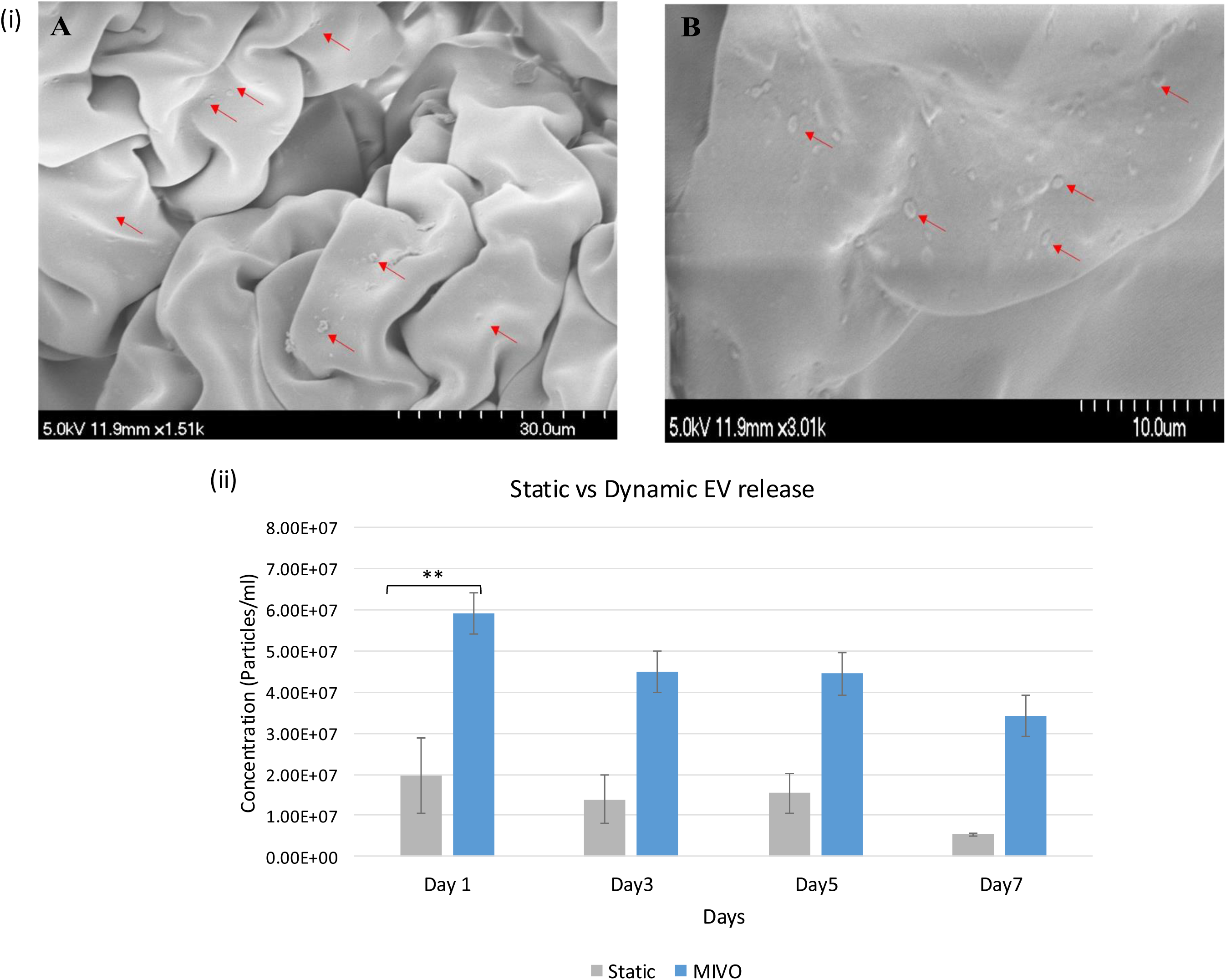
Establishing EV loaded hydrogels for sustained EV release (i) Scanning electron microscopy images of distribution of GFP-EVs loaded HA-TA hydrogels under magnifications of (A) 1500x (B) 3000x (ii) Comparison of released EV concentrations from 2%1xHA-TA hydrogels analysed by NTA overtime in Static and dynamic MIVO conditions over a period of 7 days (n=3) graph shows mean ± SEM, (**P value=0.015)

## Discussion

The therapeutic potential of sEVs lies in the ability to influence recipient cell function, but application in the clinical setting will require large quantities of purified sEV harvests with minimal co-precipitating lipoproteins. A further challenge is the high susceptibility of sEVs to rapid degradation and clearance from circulation by renal filtration and macrophage-mediated removal (21,22). This study aimed to develop HA-TA hydrogel-based matrices for the sustained release of fluorescently labelled sEVs produced in a scalable 3D dynamic bioreactor.

Multiple EV labelling and imaging techniques are currently being developed to support tracking of migration and uptake (23). While the use of lipophilic dyes was initially popular, these suffer from the potential to diffuse to adjacent cell membranes, leading to inaccurate representation of EV distribution due to the non-covalent interactions between dyes and EV surfaces (24–26). For example, Dominkus *et al.* (27) revealed the formation of PKH-26 nanoparticles that exhibited similar characteristics to PKH-26-labeled EVs in terms of size, surface area, and fluorescence intensity. To overcome this limitation, genetic manipulation strategies that involve loading reporter proteins either within the cargo or on the surface proteins have been found to ensure consistent labelling of specific EV subtypes and to minimize repetitive wash steps that could impact EV structure, concentration, and function (24,28,29). In this study, GFP-EVs were successfully generated by transducing the parent cell line with a CD-63-GFP lentiviral construct, ensuring stable GFP expression in all EV harvests expressing CD-63 protein without adverse effects on EV size and concentration. The data presented are supported by the findings of Giulia et al. (30) which demonstrated no significant differences in the proteomic profiles of GFP-tagged EVs compared to control EV samples. Additional research is warranted to explore the functional effects on ligand-receptor interactions. While not translatable to the clinical setting, this labelling approach holds immense value for EV trafficking in *in vitro* micro physiological systems and *in vivo* models of disease.

A significant limitation in GMP-grade production of EVs is the requirement for large-scale production, which in turn restricts the development of EV-based therapeutics (31). Scalability is crucial to meet the high dosage requirements in the clinical setting. This challenge encompasses not only the lack of standardized culture conditions for EV production but also the variability in EV isolation techniques (32,33). Several pharmaceutical companies specializing in EV therapeutics prefer 3D bioreactor systems, which offer closed environments to minimize contamination and produce high yields of EVs over extended periods (34). Studies using different cell types have demonstrated the consistency of EVs produced in 3D formats regarding size, concentration, and surface glycan profiles (35,36). To ensure relevance and reproducibility of initial laboratory studies aimed towards later translation, scalable bioreactor systems should ideally be used. Culturing cells in this 3D dynamic setting prevents formation of a concentration gradient of nutrients or pH that are common in static 2D cultures. The move away from use of large multiples of culture flasks is also advantageous from a space/footprint and environmental/plastics perspective. Additionally, there is the potential to employ serum free conditions once stable culture has been established, as demonstrated here. This removes the confounding serum-derived EVs. The bioreactor system supports retention of the parental cell characteristics (37). This was confirmed here, where GFP expression was retained in both cells and secreted sEVs. Use of the hollow-fiber bioreactor, coupled with SEC-based sEV isolation, resulted in high yield and sustained GFP-sEV production over time. Furthermore, replacing serum with media supplements in the culture conditions resulted in a reduction in serum-protein contaminants without negatively affecting cell viability or sEV yields. The impact of 3D culture on sEV cargo and biophysical characteristics remains poorly understood and warrants further assessment.

Among the EV-based clinical trials, 78% either did not specify the planned number of EV doses or intended to administer five or more doses. This underscores the rapid clearance of EVs from the body, necessitating high doses over extended periods (38,39). The common modes of EV administration include systemic, local, intrathecal, and intranasal delivery, which significantly impact EV efficacy due to their low circulating half-life (40,41). Therefore, this study aimed to establish tyramine modified hyaluronic acid hydrogels for sustained and prolonged EV delivery. Currently, the FDA and EMA have approved over 30 injectable hydrogel-based products for delivery of therapeutics (42). Tyramine-modified hyaluronic acid hydrogels have already demonstrated effective delivery of nanoparticles to promote delivery of angiogenic factors in a pre-clinical study targeting ischaemic myocardium (11). The study confirmed sustained delivery of star-PGA-VEGF nanoparticles incorporated into HA-TA hydrogels. The hydrogels demonstrated biocompatibility while also maintaining the bioactivity of VEGF. EVs can integrate into the extracellular matrix (ECM) (43). Thus, hydrogels used as carriers for EV loading and release must mimic the ECM to preserve EV bioactivity. Optimizing traditional factors like pore size, degradability, and compatibility can significantly enhance the *in vivo* retention of EVs (5). The novel data presented demonstrated the ability of 2% HA-TA hydrogels for sustained EV release. These hydrogels were selected based on their consistent swelling profiles and resistance to degradation over 72 hours. It has been established hydrogel swelling is a key parameter controlling cargo release from hydrogels (44). Swelling data indicated that the 2% HA-TA hydrogels may provide a more extended release profile than the 1% hydrogels, which showed signs of degradation early in the swelling experiment.

The sEVs were successfully incorporated into the HA-TA formulations, with particle dispersion observed throughout the hydrogel. It is now widely accepted that quantifying sEVs by protein content is inaccurate (41,45–48) and so EV release patterns from the hydrogels were quantified by tracking analysis. The kinetics of sEV release was investigated in a 3D dynamic model, enabling real-time analysis of sEV release at various time points. Lower concentrations of sEVs were observed in static conditions. This may be due to detection limitations with NTA, or due to sEVs remaining trapped within the hydrogels. In dynamic conditions, sEVs were released from the hydrogels consistently over seven days, indicating the potential of these 2% HA-TA hydrogels for sustaining sEV release. The findings reveal a significant enhancement in sustained sEV release from the hydrogel within the dynamic system. Another study demonstrated consistent release of miRNA 24-3p-rich EVs from di(ethylene glycol) monomethyl ether methacrylate (DEGMA)-functionalized hyaluronic acid in a rabbit model of corneal regeneration. Upon consistent treatment with the gels for 28 days, rapid healing of corneal defects and alkali burn injuries was observed (12). These preliminary studies support the potential clinical relevance of HA for EV delivery. The current study demonstrates the potential for scalable production of engineered EVs in serum free conditions and subsequent incorporation into HA-TA hydrogels for sustained release. With the establishment of a suitable hydrogel architecture, future studies employing varied EV concentrations over extended time frames are now warranted. Functional analysis of sEV released over an extended time period will also be required. The combination of sEVs and biocompatible tuneable HA-TA holds immense promise for delivery of therapeutic EVs in a variety of disease settings.

## Funding Acknowledgement

Y.C. was supported by Science Foundation Ireland and the Engineering and Physical Sciences Research Council Centre for Doctoral Training in Engineered Tissues for Discovery, Industry and Medicine (Grant numbers 18/EPSRC-CDT/3583 and EP/S02347X/1). This publication has emanated from research supported in part by a grant from Science Foundation Ireland (SFI) and the European Regional Development Fund (ERDF) under grant number 13/RC/2073_2. Additional support was received from the National Breast Cancer Research Institute (NBCRI) Ref FY24001.

## References

1. Raposo G, Stahl PD. Extracellular vesicles: a new communication paradigm? Nat Rev Mol Cell Biol. 2019 Sep;20(9):509–10.

2. Tkach M, Théry C. Communication by Extracellular Vesicles: Where We Are and Where We Need to Go. Cell. 2016 Mar 10;164(6):1226–32.

3. Chabria Y, Duffy GP, Lowery AJ, Dwyer RM. Hydrogels: 3D Drug Delivery Systems for Nanoparticles and Extracellular Vesicles. Biomedicines. 2021 Nov;9(11):1694.

4. Pinheiro A, Silva AM, Teixeira JH, Gonçalves RM, Almeida MI, Barbosa MA, et al. Extracellular vesicles: intelligent delivery strategies for therapeutic applications. Journal of Controlled Release. 2018 Nov 10;289:56–69.

5. Xie Y, Guan Q, Guo J, Chen Y, Yin Y, Han X. Hydrogels for Exosome Delivery in Biomedical Applications. Gels. 2022 Jun;8(6):328.

6. Murali VP, Holmes CA. Biomaterial-based extracellular vesicle delivery for therapeutic applications. Acta Biomaterialia. 2021 Apr 1;124:88–107.

7. Hashemi A, Ezati M, Nasr MP, Zumberg I, Provaznik V. Extracellular Vesicles and Hydrogels: An Innovative Approach to Tissue Regeneration. ACS Omega. 2024 Feb 13;9(6):6184–218.

8. Burdick JA, Prestwich GD. Hyaluronic Acid Hydrogels for Biomedical Applications. Advanced Materials. 2011;23(12):H41–56.

9. Zheng Z, Yang X, Zhang Y, Zu W, Wen M, Liu T, et al. An injectable and pH-responsive hyaluronic acid hydrogel as metformin carrier for prevention of breast cancer recurrence. Carbohydrate Polymers. 2023 Mar 15;304:120493.

10. Conte R, De Luca I, Valentino A, Cerruti P, Pedram P, Cabrera-Barjas G, et al. Hyaluronic Acid Hydrogel Containing Resveratrol-Loaded Chitosan Nanoparticles as an Adjuvant in Atopic Dermatitis Treatment. Journal of Functional Biomaterials. 2023 Feb;14(2):82.

11. O’Dwyer J, Murphy R, Dolan EB, Kovarova L, Pravda M, Velebny V, et al. Development of a nanomedicine-loaded hydrogel for sustained delivery of an angiogenic growth factor to the ischaemic myocardium. Drug Deliv Transl Res. 2020 Apr;10(2):440–54.

12. Sun X, Song W, Teng L, Huang Y, Liu J, Peng Y, et al. MiRNA 24-3p-rich exosomes functionalized DEGMA-modified hyaluronic acid hydrogels for corneal epithelial healing. Bioactive Materials. 2023 Jul 1;25:640–56.

13. Sang X, Zhao X, Yan L, Jin X, Wang X, Wang J, et al. Thermosensitive Hydrogel Loaded with Primary Chondrocyte-Derived Exosomes Promotes Cartilage Repair by Regulating Macrophage Polarization in Osteoarthritis. Tissue Eng Regen Med. 2022 Jun 1;19(3):629–42.

14. Huang L, Yang X, Deng L, Ying D, Lu A, Zhang L, et al. Biocompatible Chitin Hydrogel Incorporated with PEDOT Nanoparticles for Peripheral Nerve Repair. ACS Appl Mater Interfaces. 2021 Apr 14;13(14):16106–17.

15. Jafari D, Malih S, Eini M, Jafari R, Gholipourmalekabadi M, Sadeghizadeh M, et al. Improvement, scaling-up, and downstream analysis of exosome production. Critical Reviews in Biotechnology. 2020 Nov 16;40(8):1098–112.

16. Grangier A, Branchu J, Volatron J, Piffoux M, Gazeau F, Wilhelm C, et al. Technological advances towards extracellular vesicles mass production. Advanced Drug Delivery Reviews. 2021 Sep 1;176:113843.

17. Moloney BM, Gilligan KE, Joyce DP, O’Neill CP, O’Brien KP, Khan S, et al. Investigating the Potential and Pitfalls of EV-Encapsulated MicroRNAs as Circulating Biomarkers of Breast Cancer. Cells. 2020 Jan;9(1):141.

18. O’Brien KP, Khan S, Gilligan KE, Zafar H, Lalor P, Glynn C, et al. Employing mesenchymal stem cells to support tumor-targeted delivery of extracellular vesicle (EV)-encapsulated microRNA-379. Oncogene. 2018 Apr;37(16):2137–49.

19. Rasband W. ImageJ, U.S. National Institutes of Health, Bethesda, Maryland, USA. http://imagej.nih.gov/ij/ [Internet]. 2011 [cited 2024 Dec 2]; Available from: https://cir.nii.ac.jp/crid/1570854175774652672

20. Paciello A, Santonicola MG. Supramolecular polycationic hydrogels with high swelling capacity prepared by partial methacrylation of polyethyleneimine. RSC Adv. 2015 Oct 19;5(108):88866–75.

21. Kimiz-Gebologlu I, Oncel SS. Exosomes: Large-scale production, isolation, drug loading efficiency, and biodistribution and uptake. Journal of Controlled Release. 2022 Jul 1;347:533–43.

22. Li J, Lee Y, Johansson HJ, Mäger I, Vader P, Nordin JZ, et al. Serum-free culture alters the quantity and protein composition of neuroblastoma-derived extracellular vesicles. J Extracell Vesicles. 2015 May 27;4:10.3402/jev.v4.26883.

23. Li YJ, Wu JY, Wang JM, Hu XB, Xiang DX. Emerging strategies for labeling and tracking of extracellular vesicles. Journal of Controlled Release. 2020 Dec 10;328:141–59.

24. Gupta D, Liang X, Pavlova S, Wiklander OPB, Corso G, Zhao Y, et al. Quantification of extracellular vesicles in vitro and in vivo using sensitive bioluminescence imaging. Journal of Extracellular Vesicles. 2020;9(1):1800222.

25. Watson DC, Bayik D, Srivatsan A, Bergamaschi C, Valentin A, Niu G, et al. Efficient production and enhanced tumor delivery of engineered extracellular vesicles. Biomaterials. 2016 Oct 1;105:195–205.

26. Wen SW, Sceneay J, Lima LG, Wong CSF, Becker M, Krumeich S, et al. The Biodistribution and Immune Suppressive Effects of Breast Cancer–Derived Exosomes. Cancer Research. 2016 Nov 30;76(23):6816–27.

27. Pužar Dominkuš P, Stenovec M, Sitar S, Lasič E, Zorec R, Plemenitaš A, et al. PKH26 labeling of extracellular vesicles: Characterization and cellular internalization of contaminating PKH26 nanoparticles. Biochim Biophys Acta Biomembr. 2018 Jun;1860(6):1350–61.

28. Takahashi Y, Nishikawa M, Shinotsuka H, Matsui Y, Ohara S, Imai T, et al. Visualization and in vivo tracking of the exosomes of murine melanoma B16-BL6 cells in mice after intravenous injection. Journal of Biotechnology. 2013 May;165(2):77–84.

29. Morishita M, Takahashi Y, Nishikawa M, Sano K, Kato K, Yamashita T, et al. Quantitative analysis of tissue distribution of the B16BL6-derived exosomes using a streptavidin-lactadherin fusion protein and iodine-125-labeled biotin derivative after intravenous injection in mice. J Pharm Sci. 2015 Feb;104(2):705–13.

30. Corso G, Heusermann W, Trojer D, Görgens A, Steib E, Voshol J, et al. Systematic characterization of extracellular vesicle sorting domains and quantification at the single molecule – single vesicle level by fluorescence correlation spectroscopy and single particle imaging. J Extracell Vesicles. 2019 Sep 18;8(1):1663043.

31. Rezaie J, Feghhi M, Etemadi T. A review on exosomes application in clinical trials: perspective, questions, and challenges. Cell Communication and Signaling. 2022 Sep 19;20(1):145.

32. Jakl V, Ehmele M, Winkelmann M, Ehrenberg S, Eiseler T, Friemert B, et al. A novel approach for large-scale manufacturing of small extracellular vesicles from bone marrow-derived mesenchymal stromal cells using a hollow fiber bioreactor. Frontiers in Bioengineering and Biotechnology [Internet]. 2023 [cited 2023 Oct 24];11. Available from: https://www.frontiersin.org/articles/10.3389/fbioe.2023.1107055

33. Paolini L, Monguió-Tortajada M, Costa M, Antenucci F, Barilani M, Clos-Sansalvador M, et al. Large-scale production of extracellular vesicles: Report on the “massivEVs” ISEV workshop. Journal of Extracellular Biology. 2022;1(10):e63.

34. Rohde E, Pachler K, Gimona M. Manufacturing and characterization of extracellular vesicles from umbilical cord–derived mesenchymal stromal cells for clinical testing. Cytotherapy. 2019 Jun 1;21(6):581–92.

35. Ghodasara A, Raza A, Wolfram J, Salomon C, Popat A. Clinical Translation of Extracellular Vesicles. Advanced Healthcare Materials. n/a(n/a):2301010.

36. Gobin J, Muradia G, Mehic J, Westwood C, Couvrette L, Stalker A, et al. Hollow-fiber bioreactor production of extracellular vesicles from human bone marrow mesenchymal stromal cells yields nanovesicles that mirrors the immuno-modulatory antigenic signature of the producer cell. Stem Cell Research & Therapy. 2021 Feb 12;12(1):127.

37. Syromiatnikova V, Prokopeva A, Gomzikova M. Methods of the Large-Scale Production of Extracellular Vesicles. Int J Mol Sci. 2022 Sep 10;23(18):10522.

38. Duong A, Parmar G, Kirkham AM, Burger D, Allan DS. Registered clinical trials investigating treatment with cell-derived extracellular vesicles: a scoping review. Cytotherapy. 2023 Sep;25(9):939–45.

39. M.D. Anderson Cancer Center. Phase I Study of Mesenchymal Stromal Cells-Derived Exosomes With KrasG12D siRNA for Metastatic Pancreas Cancer Patients Harboring KrasG12D Mutation [Internet]. clinicaltrials.gov; 2021 Apr [cited 2021 May 19]. Report No.: NCT03608631. Available from: https://clinicaltrials.gov/ct2/show/NCT03608631

40. Johnson J, Law SQK, Shojaee M, Hall AS, Bhuiyan S, Lim MBL, et al. First-in-human clinical trial of allogeneic, platelet-derived extracellular vesicles as a potential therapeutic for delayed wound healing. Journal of Extracellular Vesicles. 2023;12(7):12332.

41. Lu W, Zeng M, Liu W, Ma T, Fan X, Li H, et al. Human urine-derived stem cell exosomes delivered via injectable GelMA templated hydrogel accelerate bone regeneration. Materials Today Bio. 2023 Apr 1;19:100569.

42. Mandal A, Clegg JR, Anselmo AC, Mitragotri S. Hydrogels in the clinic. Bioengineering & Translational Medicine. 2020;5(2):e10158.

43. Patel NJ, Ashraf A, Chung EJ. Extracellular Vesicles as Regulators of the Extracellular Matrix. Bioengineering (Basel). 2023 Jan 19;10(2):136.

44. Bettini R, Colombo P, Massimo G, Catellani PL, Vitali T. Swelling and drug release in hydrogel matrices: polymer viscosity and matrix porosity effects. European Journal of Pharmaceutical Sciences. 1994 Oct 1;2(3):213–9.

45. Théry C, Witwer KW, Aikawa E, Alcaraz MJ, Anderson JD, Andriantsitohaina R, et al. Minimal information for studies of extracellular vesicles 2018 (MISEV2018): a position statement of the International Society for Extracellular Vesicles and update of the MISEV2014 guidelines. Journal of Extracellular Vesicles. 2018 Dec 1;7(1):1535750.

46. Liu X, Wu C, Zhang Y, Chen S, Ding J, Chen Z, et al. Hyaluronan-based hydrogel integrating exosomes for traumatic brain injury repair by promoting angiogenesis and neurogenesis. Carbohydrate Polymers. 2023 Apr 15;306:120578.

47. Born LJ, McLoughlin ST, Dutta D, Mahadik B, Jia X, Fisher JP, et al. Sustained Released of Bioactive Mesenchymal Stromal Cell-Derived Extracellular Vesicles from 3D-Printed Gelatin Methacrylate Hydrogels [Internet]. bioRxiv; 2021 [cited 2023 Oct 18]. p. 2021.09.28.462252. Available from: https://www.biorxiv.org/content/10.1101/2021.09.28.462252v1

48. Welsh JA, Goberdhan DCI, O’Driscoll L, Buzas EI, Blenkiron C, Bussolati B, et al. Minimal information for studies of extracellular vesicles (MISEV2023): From basic to advanced approaches. Journal of Extracellular Vesicles. 2024;13(2):e12404.

