## Supplementary Figure 1 for "Hyaluronic acid hydrogels: Establishing a sustained delivery system for extracellular vesicles"

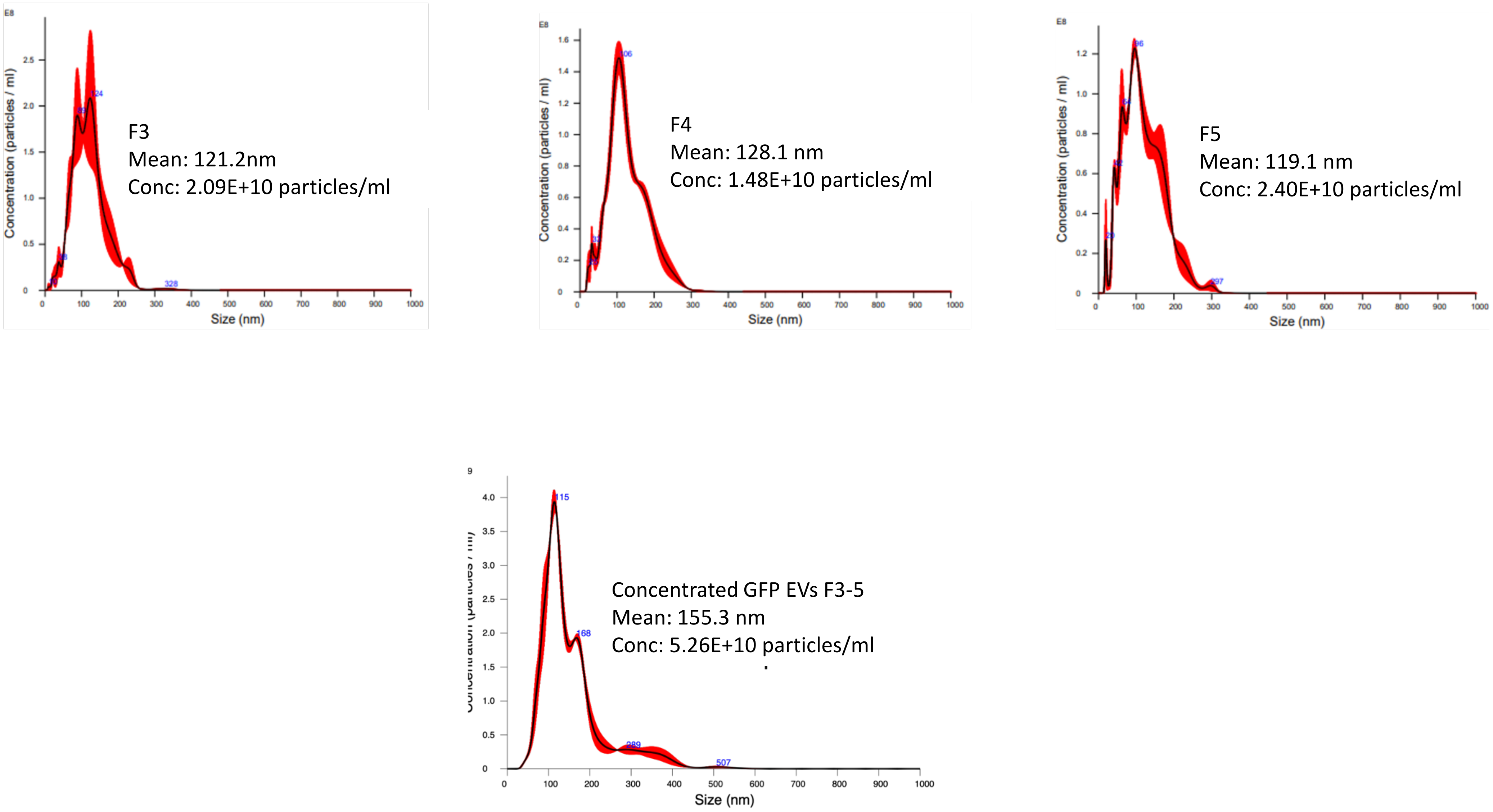


**Supplementary Figure 1**: Representative NTA graphs demonstrating size distribution and EV concentrations of concentrated GFP-EV fractions obtained by concentrating fractions (F3-F5) by ultrafiltration in triplicates, (n=3)
