## Supplementary Table 1 for "Hyaluronic acid hydrogels: Establishing a sustained delivery system for extracellular vesicles"

1%1x PBS

| 1 hr | 2 hrs | 3 hrs | 4 hrs | 5 hrs | 24 hrs | 48 hrs | 72 hrs |
|---|---|---|---|---|---|---|---|
| -5.78 | -10.35 | -11.49 | -12.10 | -12.48 | -12.56 | -14.16 | -19.94 |
| -19.63 | -22.98 | -26.71 | -25.27 | -27.32 | -27.70 | -24.96 | -26.03 |
| 11.87 | 5.71 | 5.71 | 6.47 | 2.97 | 2.21 | 2.74 | 6.85 |

1%1x Media

| 1 hr | 2 hrs | 3 hrs | 4 hrs | 5 hrs | 24 hrs | 48 hrs | 72 hrs |
|---|---|---|---|---|---|---|---|
| -3.71 | -5.01 | -5.34 | -5.86 | -5.99 | -6.18 | -7.55 | -8.53 |
| -35.16 | -38.02 | -36.00 | -37.17 | -36.98 | -36.20 | -39.00 | -35.74 |
| -22.40 | -24.61 | -25.20 | -24.80 | -25.78 | -24.54 | -24.15 | -26.63 |

1%2x PBS

| 1 hr | 2 hrs | 3 hrs | 4 hrs | 5 hrs | 24 hrs | 48 hrs | 72 hrs |
|---|---|---|---|---|---|---|---|
| -8.46 | -12.31 | -13.92 | -12.00 | -5.54 | -10.92 | -13.08 | -13.85 |
| -8.49 | -13.12 | 7.65 | 10.78 | 5.06 | 6.74 | 6.68 | 1.75 |
| -6.93 | -7.09 | -12.43 | -12.83 | -14.02 | -13.63 | -15.14 | -14.10 |

1%2x PBS

| 1 hr | 2 hrs | 3 hrs | 4 hrs | 5 hrs | 24 hrs | 48 hrs | 72 hrs |
|---|---|---|---|---|---|---|---|
| -7.18 | -7.25 | -9.80 | -9.95 | -5.56 | -6.40 | -8.64 | -10.19 |
| -6.04 | -27.24 | -25.31 | -24.85 | -25.93 | -25.39 | -26.62 | -24.15 |
| -6.39 | -0.77 | -2.62 | -2.01 | -3.40 | -3.86 | -2.70 | -0.93 |

**Supplementary Table 1**: Weight measurements of blank 1% (1x and 2x) hydrogels observed at 1-5 hours upto 72 hours to assess swelling behaviour patterns. Reduction in initial weight ($m_0$) measured was observed in all 1% HA-TA hydrogels
